# Disruption of undecaprenyl phosphate recycling suppresses *ampC* beta-lactamase induction in *Pseudomonas aeruginosa*

**DOI:** 10.1101/2025.06.03.657773

**Authors:** Karina Klycheva, Joël Gyger, Mélissa Frund, Gabriel Torrens, Felipe Cava, Coralie Fumeaux

## Abstract

Beta-lactam antibiotics are widely used to treat bacterial infections, but their efficacy is compromised by resistance mechanisms such as the production of beta-lactamases. In *Pseudomonas aeruginosa*, the chromosomally encoded beta-lactamase AmpC is the primary mediator of beta-lactam resistance. *ampC* expression is regulated by the transcription factor AmpR, which responds to intracellular peptidoglycan (PG) fragments. Under normal conditions, AmpR binds the PG precursor (UDP-MurNAc-pentapeptide, UDP-M5) and represses *ampC* expression. However, during beta-lactam treatment or in PG recycling-deficient mutants such as *ampD* mutants, PG degradation products (anhydromuropeptides, AMP) accumulate and activate AmpR, resulting in elevated *ampC* expression and beta-lactam resistance.

We hypothesized that shifting the balance of PG precursors could modulate AmpR activity and suppress beta-lactamase expression, even in derepressed strains. Undecaprenyl phosphate (UndP) is a lipid carrier essential for translocating PG precursors across the bacterial inner membrane. Recent work has identified members of the DedA superfamily as UndP flippases responsible for recycling this lipid carrier. Disruption of UndP recycling leads to cytoplasmic accumulation of UDP-M5, the known AmpR repressor. Here, we show that deletion of *dedA4*, which encodes a predicted UndP flippase in *P. aeruginosa*, causes UDP-M5 accumulation and significantly reduces AmpC production and beta-lactam resistance in an *ampD* mutant. These findings highlight the influence of PG precursor dynamics on beta-lactamase regulation and identify DedA4 as a promising therapeutic target. Inhibiting UndP recycling offers a novel strategy to counteract beta-lactam resistance in *P. aeruginosa* and potentially other AmpC-producing pathogens.

**IMPORTANCE:** Beta-lactam resistance remains a major clinical challenge, often driven by the overexpression of chromosomally encoded beta-lactamases such as AmpC in *Pseudomonas aeruginosa*. Strategies to limit *ampC* activation are urgently needed to preserve the efficacy of these frontline antibiotics. Our study identifies DedA4, a membrane protein involved in undecaprenyl phosphate (UndP) recycling, as a potential target for such intervention. Inactivation of DedA4 disrupts UndP transport, leading to the accumulation of soluble, membrane-unbound PG precursors which act as corepressors of AmpC expression. This disruption in the balance between cell wall synthesis and turnover restores susceptibility to β-lactams, even in strains harboring clinically significant resistance mutations such as those in *ampD* or *dacB*. Targeting UndP flippases like DedA4 could therefore represent a novel adjuvant strategy to combat AmpC-mediated beta-lactam resistance.

## INTRODUCTION

*Pseudomonas aeruginosa* is an opportunistic pathogen that poses a significant clinical challenge due to its intrinsic and acquired mechanisms of antibiotic resistance (1, 2). A major contributor to this resistance is the chromosomally encoded AmpC beta-lactamase, which can hydrolyze a broad range of beta-lactam antibiotics, reducing their therapeutic efficacy (3–5). The expression of the *ampC* gene is tightly regulated and responsive to cell wall stress, particularly during exposure to certain beta-lactams (5). Under normal conditions, *ampC* is expressed at low basal levels. However, treatment with beta-lactamase-inducing antibiotics, such as cefoxitin and imipenem, strongly upregulates *ampC*, whereas non-inducers like piperacillin and ceftazidime typically do not (3, 4). Non-inducing beta-lactams retain activity against *P. aeruginosa*, but become ineffective when *ampC* is dysregulated (4, 6).

Mutations in cell wall-related genes, such as *ampD* or *dacB*, can lead to constitutive overexpression of *ampC*, resulting in high-level resistance, even to non-inducing beta-lactams. These mutants are clinically relevant, having been isolated from patients and associated with therapeutic failure (5–9). Consequently, there is growing interest in dissecting the molecular pathways that govern *ampC* expression and identifying strategies to counteract its upregulation.

The transcriptional regulator AmpR controls *ampC* expression in response to the balance between specific peptidoglycan (PG) biosynthetic precursors and recycling products (5, 7–9). In wild-type cells, PG turnover fragments are efficiently recycled and AmpR binds cytoplasmic PG precursors (UDP-M5), forming a tetrameric complex that represses *ampC* transcription (10, 11) (**Fig. 1A)**. In contrast, during a beta-lactam treatment or in *ampD* or *dacB* mutants, an abnormal buildup of PG recycling intermediates (anhydromuropeptides, AMP), switches AmpR into its activator state, leading to constitutive high-level *ampC* expression (5, 12, 13) (**Fig. 1B**).

**Figure 1.**
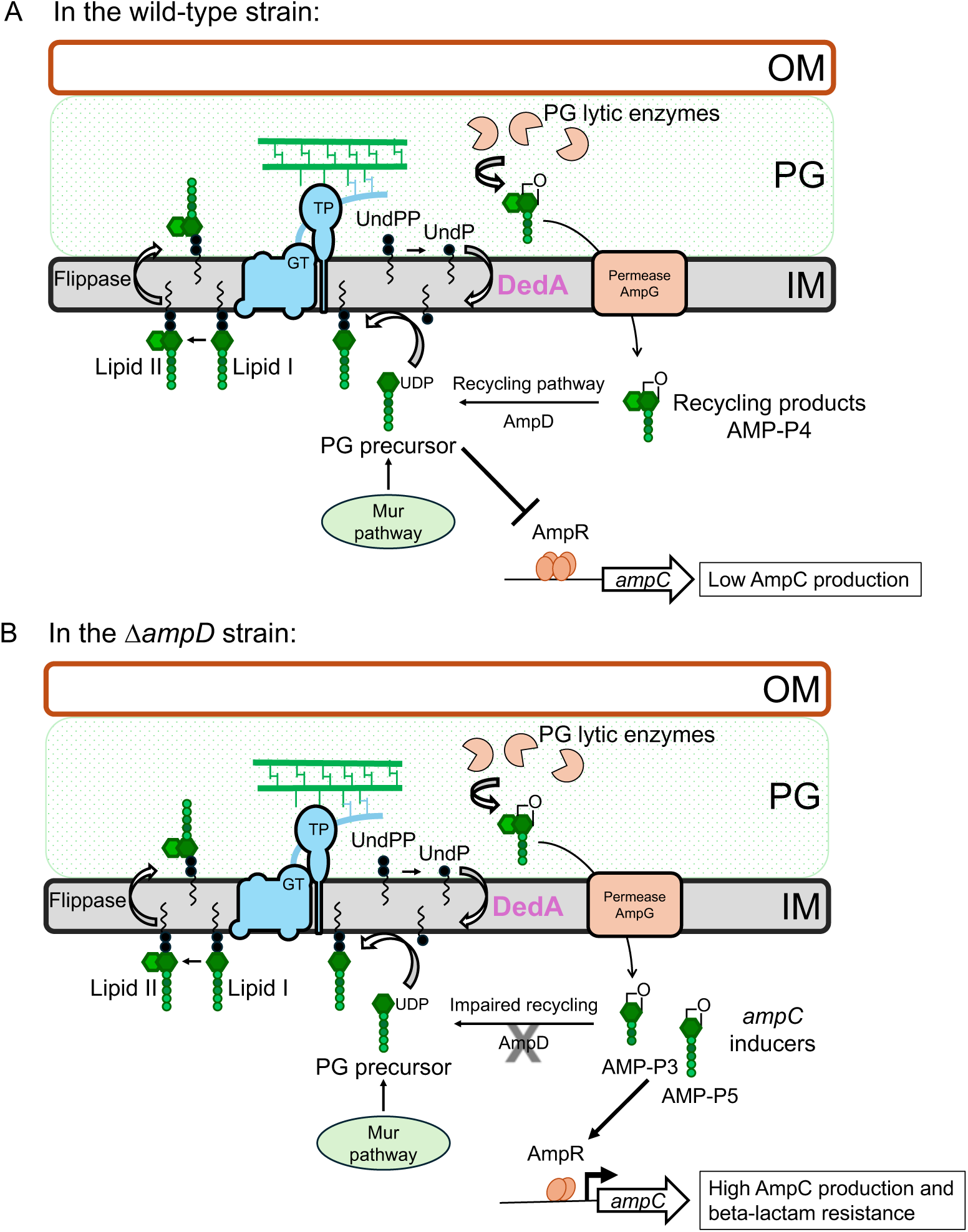
Overview of peptidoglycan (PG) cell wall synthesis, undecaprenyl phosphate (UndP) cycle and *ampC* regulation in *P. aeruginosa* wild-type and Δ*ampD* strains. (**A**) Peptidoglycan precursors are synthesized in the cytoplasm via the Mur pathway and loaded onto the lipid carrier undecaprenyl phosphate (UndP) (14–16). These UndP-linked precursors are transported across the membrane and incorporated into the expanding PG matrix by glycosyltransferase (GT) and transpeptidase (TP) enzymes (16). Undecaprenyl pyrophosphate (UndPP), generated in the process, is dephosphorylated and flipped by members of the DedA superfamily of flippases to regenerate cytoplasmic-facing UndP (18, 19). Mature PG is degraded by PG lytic enzymes to produce anhydro-MurNAc-containing muropeptide (AMP) turnover products, which are imported into the cytoplasm by the AmpG permease for recycling (5, 52). Tetrapeptide-AMP products (AMP-P4) are rapidly processed by recycling enzymes to form new PG precursors. These precursors prevent the activation of *ampC* expression by AmpR, resulting in *ampC* repression in the absence of cell wall damage (9, 10). (**B**) In the *ampD* mutant, PG recycling is impaired, leading to the accumulation of tripeptide-AMP (AMP-P3), predominantly, and to a lesser extent pentapeptide-AMP (AMP-P5) in the cytoplasm (see Fig. 6). These turnover products are thought to be potent activators of AmpR, resulting in *ampC* induction and beta-lactam resistance (8, 9, 12, 13).

We hypothesized that shifting the balance back toward PG precursors, even in derepressed mutants, could suppress *ampC* induction. One potential way to achieve this is by limiting the availability of the lipid carrier undecaprenyl phosphate (UndP), which is essential for PG synthesis (14, 15). In the cytoplasm, PG precursors are synthesized by the MurA-MurF enzymes to form UDP-MurNAc-pentapeptide (UDP-M5). This molecule is transferred to UndP by MraY to form Lipid I, which is then converted into Lipid II by MurG through addition of the second sugar UDP-GlcNAc. Lipid II is flipped across the inner membrane by the flippase MurJ and incorporated into the cell wall by glycosyltransferase and transpeptidase enzymes (16, 17). Once utilized, Lipid II generates undecaprenyl pyrophosphate (UndPP), which undergoes dephosphorylation and recycling back into UndP, ensuring continued PG synthesis (14, 15, 18–20) (**Fig. 1**).

Recent studies in *Vibrio cholerae* and *Staphylococcus aureus* have shown that deletion of UndP recycling enzymes, members of the DedA superfamily and DUF368 domain-containing proteins, leads to the cytoplasmic accumulation of UDP-M5 (18). This occurs due to an insufficient pool of UndP, despite ongoing synthesis by the *de novo* pathway via UppS (also named IspU in *Escherichia coli*) (21). The resulting accumulation of PG precursors may affect cell wall homeostasis and regulatory pathways, including *ampC* induction.

To test whether UndP recycling affects *ampC* regulation in *P. aeruginosa*, we investigated the impact of deleting its predicted UndP flippase genes. Here, we report the effects of *dedA4* deletion in both wild-type and Δ*ampD* backgrounds. We show that the disruption of UndP recycling leads to UDP-M5 accumulation and modulates AmpR activity. In *ampC*-derepressed strains, this shift in precursor balance suppresses AmpC production and beta-lactam resistance. Our findings suggest that UndP recycling is a regulatory node in beta-lactam resistance. Inhibiting UndP recycling could represent a novel therapeutic strategy to enhance beta-lactam efficacy against resistant *P. aeruginosa* strains and other AmpC-producing species, like *Citrobacter freundii* and *Enterobacter cloacae* (4).

## RESULTS

### Inactivation of P. aeruginosa dedA4 reduces AmpC production and beta-lactam resistance

As described above (**Fig. 1**), AmpR activity is influenced by cytoplasmic PG precursor levels. Based on this, we hypothesized that disrupting UndP recycling would impact PG precursor levels and reduce *ampC* induction. Unlike some other organisms, *P. aeruginosa* lacks DUF368 homologues but encodes five DedA family members (22). In *Bacillus subtilis*, the *uptA ykoX* double mutant lacks known UndP flippases, providing a system to test candidate genes for this function (19). Among the *P. aeruginosa* DedA proteins, DedA4 (PA4029) was previously shown to complement this *Bacillus* mutant, suggesting that it acts as a bona fide UndP flippase (19). Therefore, we deleted *dedA4* (*PA4029*) in wild-type and Δ*ampD P. aeruginosa* strains and assessed the effects on AmpC levels and beta-lactam resistance.

To monitor the effect of DedA4 inactivation on *ampC* induction, we tested the beta-lactam resistance of mutant strains. Consistent with prior results, the Δ*ampD* strain was highly resistant to the antipseudomonal beta-lactams, ceftazidime and piperacillin (**Fig. 2A**). Deletion of *dedA4* in the Δ*ampD* strain significantly reduced beta-lactam resistance (**Fig. 2A**), while it did not affect viability in the absence of the antibiotic. To quantify this change in resistance, we measured the minimum inhibitory concentrations (MICs) of the relevant strains. We confirmed that the double mutant Δ*ampD* Δ*dedA4* was 4 times more sensitive to ceftazidime and piperacillin than the parental Δ*ampD* strain, with MICs of 20 µg/ml (Δ*ampD*) and 5 µg/ml (Δ*ampD* Δ*dedA4*) for piperacillin, and 5 µg/ml (Δ*ampD*) and 1.25 µg/ml (Δ*ampD* Δ*dedA4*) for ceftazidime (**Table 1)**. As controls, we measured MICs for the non-beta-lactam antibiotics A22, tobramycin and gentamicin, which showed no significant changes across strains (**Table 1**). These results suggest that deletion of *dedA4* impacts specifically resistance to beta-lactam antibiotics but not cell envelope integrity.

**Figure 2.**
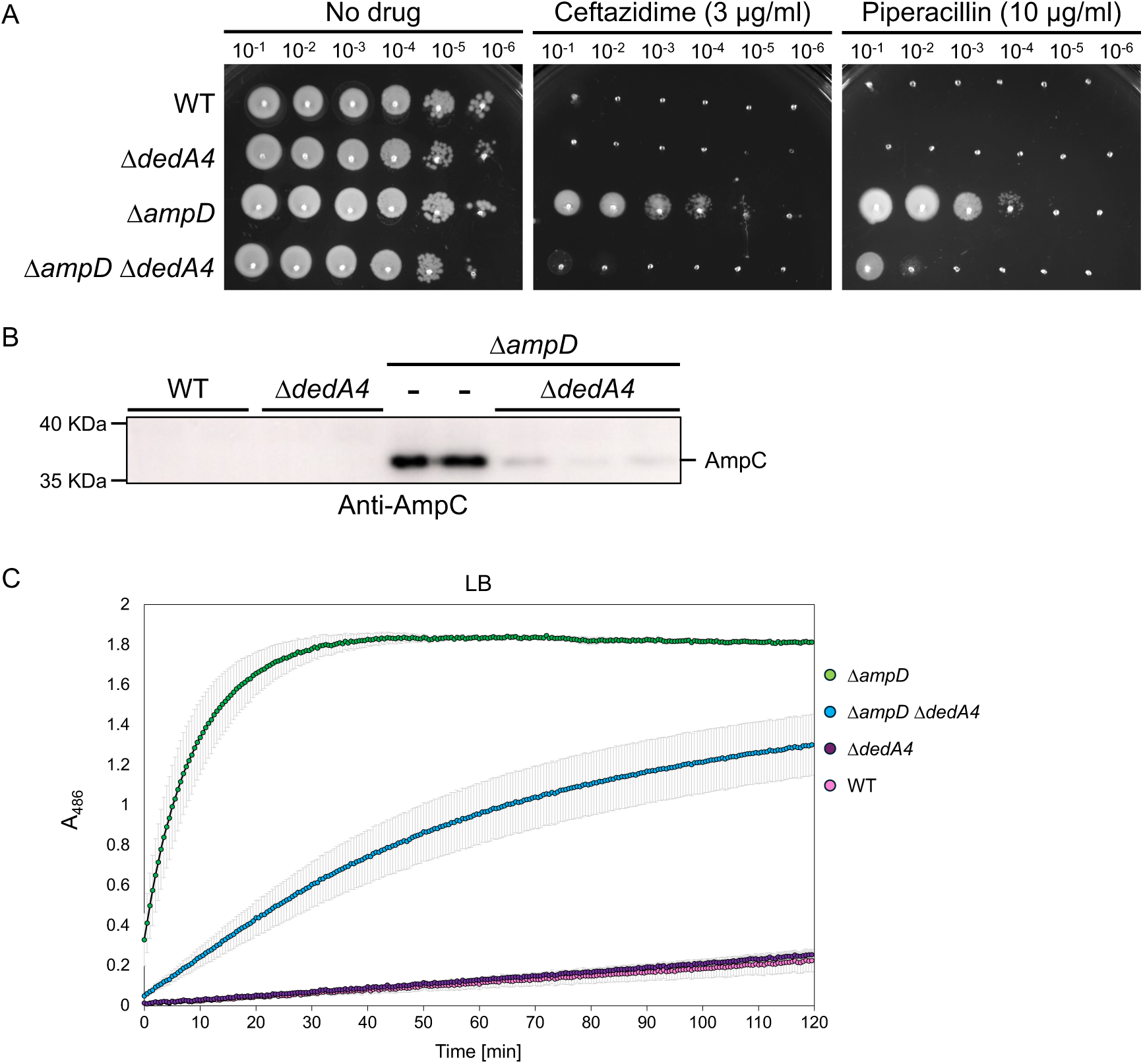
DedA4 is required for AmpC production and beta-lactam resistance in cells defective for AmpD. **(A)** Cultures of strains PAO1 [WT], CF1842 [Δ*dedA4*], CF5 [Δ*ampD*] and CF1844 [Δ*ampD* Δ*dedA4*] were serially diluted and 5 µl of each dilution was spotted onto LB agar supplemented with either ceftazidime (3 µg/ml) or piperacillin (10 µg/ml), as indicated. Plates were incubated overnight at 30°C and photographed. **(B)** Immunoblot for AmpC protein using the strains from panel **(A)**, including biological replicates. **(C)** Nitrocefin hydrolysis assays using cell lysates of the strains from panels **(A)** and **(B)**. Data represent the mean of three independent assays with two to four biological replicates; error bars indicate standard error.

**Table 1.**
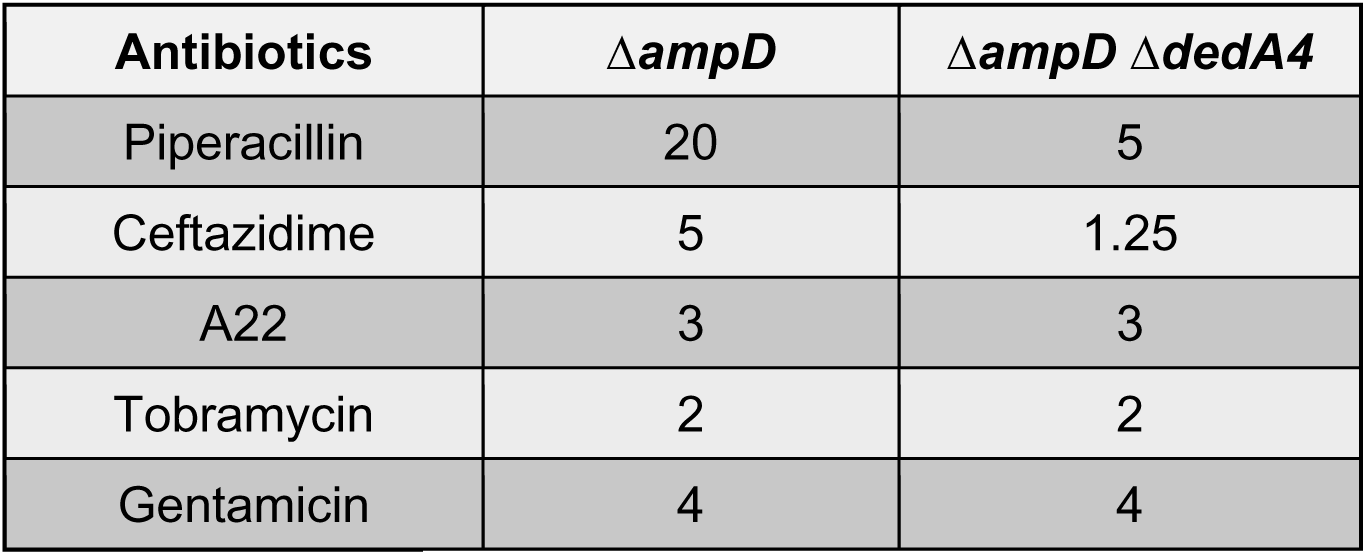
Minimal inhibitory concentration (MIC) of selected antibiotics:

| **Antibiotics** | **$\Delta ampD$** | **$\Delta ampD \Delta dedA4$** |
| --- | --- | --- |
| Piperacillin | 20 | 5 |
| Ceftazidime | 5 | 1.25 |
| A22 | 3 | 3 |
| Tobramycin | 2 | 2 |
| Gentamicin | 4 | 4 |

To determine whether DedA4 influences beta-lactam resistance through AmpC, we quantified AmpC production and activity using Western blot and nitrocefin hydrolysis assays. As expected, the production of AmpC was undetectable in the wild-type, and Δ*dedA4* strains, while it strongly accumulated in the Δ*ampD* strains. However, the level produced was clearly reduced in the double mutants Δ*ampD* Δ*dedA4* (**Fig. 2B**). Similar observations were made for AmpC activity. The level of nitrocefin hydrolysis detected for the Δ*ampD* lysates was high, with all the nitrocefin being hydrolyzed after 30 minutes of incubation, while the double mutant lysates still haven’t processed all the nitrocefin after 2 hours of incubation (**Fig. 2C**). Notably, beta-lactam resistance and AmpC production were restored back to the level of the Δ*ampD* strain upon expression of *dedA4* from a plasmid in Δ*ampD* Δ*dedA4* cells (**Fig. 3A and 3B**), indicating that the phenotype of the deletion allele was caused by the inactivation of DedA4 and not a polar effect of the deletion on the expression of the nearby genes. Expression of *ampD* in Δ*ampD* Δ*dedA4* cells from the same plasmid reversed the derepression, restoring beta-lactam sensitivity and eliminating AmpC production (**Fig. 3A and 3B**). Collectively, these results support the idea that DedA4 contributes to beta-lactam resistance by modulating *ampC* induction, likely through its role in UndP recycling.

**Figure 3.**
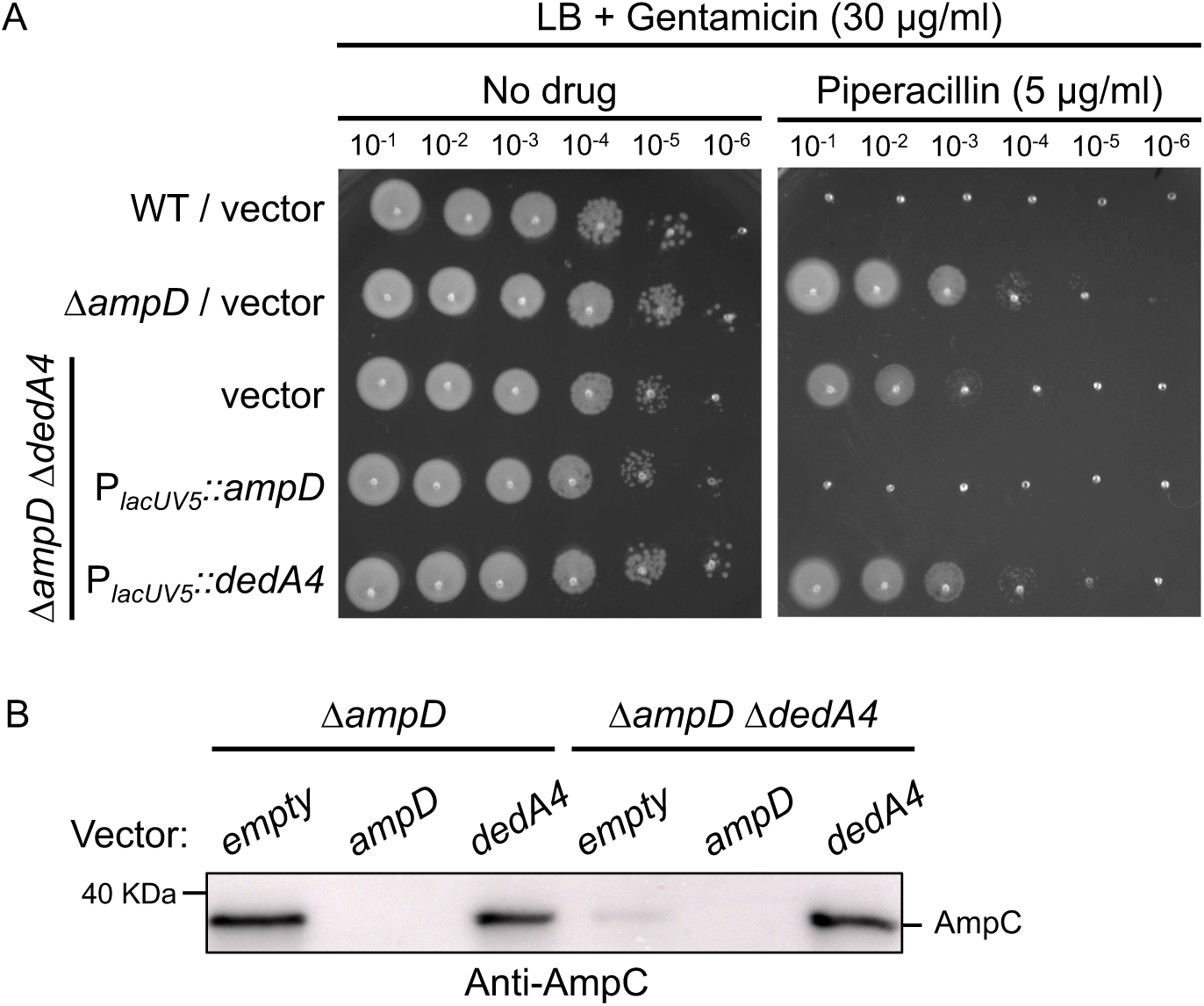
Expression of *dedA4* from a replicative plasmid restores AmpC production ad beta-lactam resistance in Δ*ampD* Δ*dedA4* cells. (**A**) Cultures of strains PAO1 [WT], CF5 [Δ*ampD*] and CF1844 [Δ*ampD* Δ*dedA4*] containing plasmids pPSV38 [vector control], pCF1098 [P*_lacUV5_*::*ampD*], or pCF835 [P*_lacUV5_*::*dedA4*] were serially diluted and 5 µl of each dilution was spotted onto LB agar supplemented with gentamicin (30 µg/ml) for plasmid maintenance, with or without piperacillin (5 µg/ml), as indicated. Plates were incubated overnight at 30°C and photographed. **(B)** Immunoblot for AmpC protein using the strains from panel **(A)**.

### UndP flippase activity is required for full ampC induction and beta-lactam resistance in *ΔampD mutants*

It was previously demonstrated that DedA4 was able to complement an *uptA ykoX* double mutant in *B. subtilis* that lacks both genes encoding UndP flippases (19). Alignment of *B. subtilis* UptA (YngC), *E. coli* YqjA and YghB and *P. aeruginosa* DedA4 and DedA5 highlighted the presence of conserved and essential residues for their function. DedA4*^Pae^* Glu38 and Asp50 likely bind protons, while Arg129 and Arg135 likely bind the negatively charged phosphate group of UndP (**Fig. 4A**) (19, 23). To test if UndP transport activity was necessary for *ampC* induction in the Δ*ampD* strain, we produced a FLAG-tagged variant of either DedA4(WT) or alleles mutated for two of these key residues. Δ*ampD* Δ*dedA4* cells expressing the functional mutants DedA4(D50A), DedA4(R129A) and the double mutant DedA4(D50A, R129A), failed to restore beta-lactam resistance and AmpC production, unlike cells expressing the WT DedA allele (**Fig. 4B and 4C**). These results indicate that DedA4 function as an UndP flippase is required for optimal *ampC* induction in the Δ*ampD* strain. All mutant variants were produced at similar levels upon IPTG (1mM) treatment (**Fig. 4D**).

**Figure 4.**
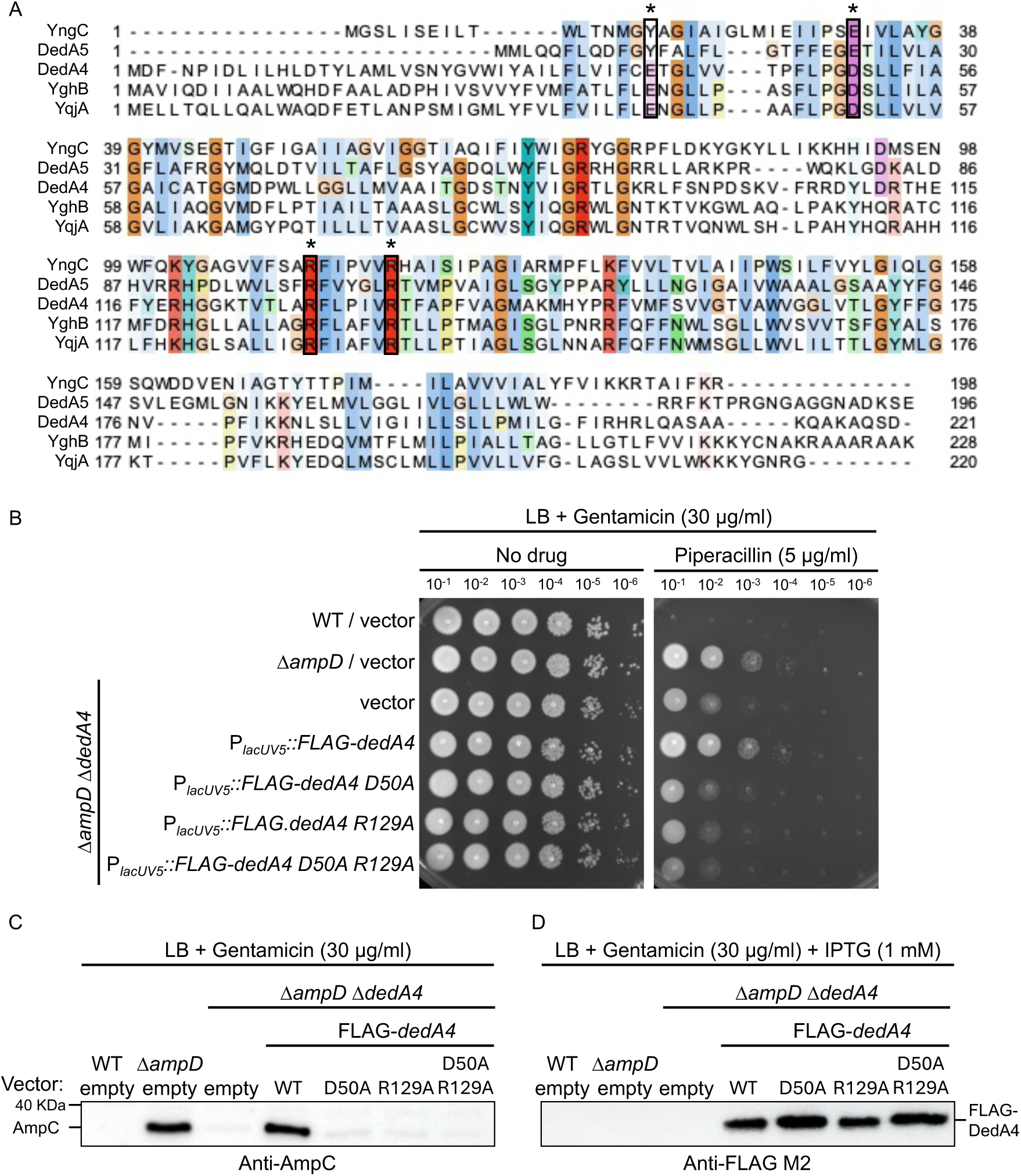
UndP recycling activity of DedA4 is required for complementation of Δ*ampD* Δ*dedA4* mutant. (**A**) Multiple sequence alignment of DedA homologues (*B. subtilis* YngC; *P. aeruginosa* DedA4 and DedA5; *E. coli* YghB and YqjA). Four highly conserved residues are boxed in black and marked with an asterisk (see text for details). Cultures of strains PAO1 [WT], CF5 [Δ*ampD*] and CF1844 [Δ*ampD* Δ*dedA4*] carrying plasmids pPSV38 [vector control], pCF661 [P*_lacUV5_*::FLAG-*dedA4*], pCF580 [P*_lacUV5_*::FLAG-*dedA4 D50A*], pCF584 [P*_lacUV5_*::FLAG-*dedA4 R129A*] or pCF1154 [P*_lacUV5_*::FLAG-*dedA4 D50A R129A*] were serially diluted and 5 µl of each dilution was spotted onto LB agar supplemented with gentamicin (30 µg/ml) for plasmid maintenance, with or without piperacillin (5 µg/ml), as indicated. (**B**) Immunoblot for AmpC protein using the strains from panel (**A**). (**C**) Immunoblot for FLAG-tagged DedA4 variants using the same strains as in panel (**A**) grown for four hours in LB supplemented with gentamicin (30 µg/ml) and IPTG (1 mM).

**Figure 5.**
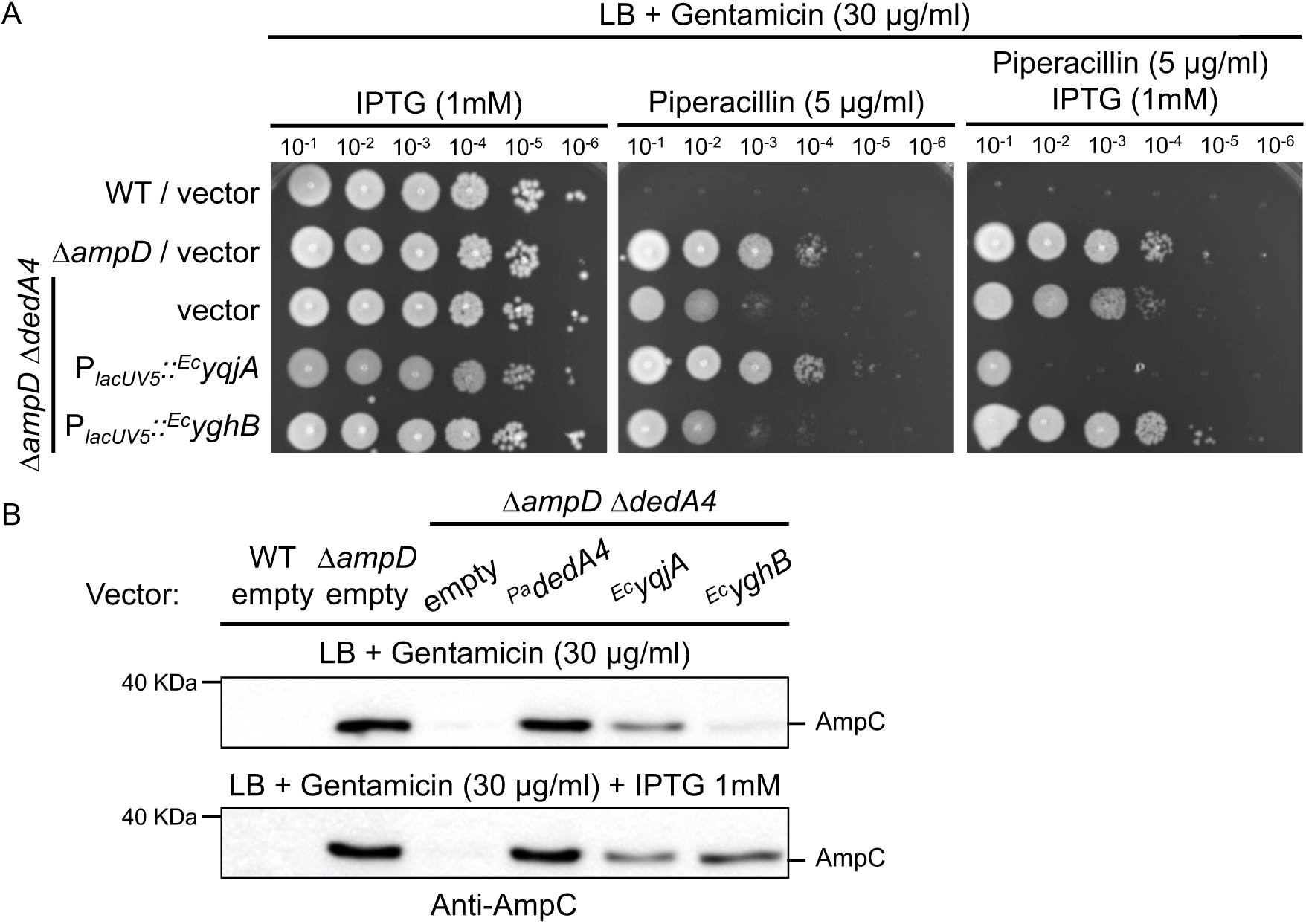
Heterologous complementation with DedA homologues from *E. coli* can complement Δ*ampD* Δ*dedA4*. (A) Cultures of strains PAO1 [WT], CF5 [Δ*ampD*] and CF1844 [Δ*ampD* Δ*dedA4*] carrying plasmids pPSV38 [vector control], pCF1147 [P*_lacUV5_*::*^Ec^yqjA*] and pCF1150 [P*_lacUV5_*::*^Ec^yghB*] were serially diluted and 5 µl of each dilution was spotted onto LB agar supplemented with gentamicin (30 µg/ml) for plasmid maintenance, with or without piperacillin (5 µg/ml) or IPTG (1 mM), as indicated. (**B**) Immunoblot for AmpC protein using the strains from panel (**A**) grown in LB supplemented with gentamicin (30 µg/ml) and with or without IPTG (1 mM), as indicated.

Since *P. aeruginosa* genome encodes five DedA paralogues (22), we attempted to complement the Δ*ampD* Δ*dedA4* mutant with each of them. Complementation only worked with *dedA4* and *dedA5* when IPTG was added (**Fig. S1A and S1B**). These results suggest that DedA4 is the major DedA family member involved in UndP recycling under our conditions. Although *dedA5* deletion in the Δ*ampD* strain had minimal impact on beta-lactam resistance (**Fig. S2**), overexpression of *dedA5* with IPTG restored AmpC production in the Δ*ampD* Δ*dedA4* background (**Fig. S1B**). This suggests that DedA5 may primarily transport a different anionic lipid or may function under distinct conditions but can substitute for UndP flippase activity when overexpressed, similar to YkoX in *B. subtilis* (19). Interestingly, overexpression of *dedA4* appears detrimental to the Δ*ampD* Δ*dedA4* strain. This is evidenced by the formation of smaller colonies on IPTG-containing plates (**Fig. S1A**) and complete growth inhibition in the presence of both IPTG and piperacillin (**Fig. S1A, right panel**), despite continued high-level AmpC production (**Fig. S1B, bottom panel**). These findings suggest that overexpression of DedA4 may have toxic effects independent of *ampC* regulation. One possibility is that excess DedA4 perturbs membrane homeostasis by acting on lipids other than UndP. Supporting this idea, other DedA family members have been implicated in the transport of diverse lipids. For example, *B. subtilis* PetA was shown to transport phosphatidylethanolamine (24).

Heterologous complementation with the UndP transporters of *E. coli* YqjA and YghB (homologues of DedA proteins), restore complete AmpC production and beta-lactam resistance to the Δ*ampD* Δ*dedA4* strain. We note that overexpression of *^Ec^yghB* with IPTG was required, like what we observed for *^Pa^dedA5*. This finding suggests that functional UndP flippase activity, regardless of its origin, is sufficient to restore *ampC* regulation in *P. aeruginosa*. We hypothesize that enough UndP lipid carriers on the inner leaflet of the cytoplasmic membrane is required to form lipid I and therefore reduced the pool of soluble PG precursors in the cytoplasm. In the absence of UndP recycling, soluble PG precursors may accumulate in the cytoplasm, bind to AmpR and maintain it in a repressive state, thereby limiting *ampC* induction.

### Disruption of UndP recycling alters the ratio of PG recycling products to PG precursors and suppresses ampC induction

The results presented thus far suggest that mutants lacking DedA4 are impaired in UndP recycling and accumulate cytoplasmic PG precursors (UDP-MurNAc-pentapeptide, UDP-M5). To test this hypothesis, we first performed muropeptide analysis to determine whether the overall structure of PG was altered in cells lacking *dedA4*. Consistent with prior results obtained in *V. cholerae* lacking *vca0040*, the *dedA4* mutant derivatives had 10% to 20% less PG than the wild-type and minor cross-linking defects (**Fig. S3A-S3C**) (18). We then quantified both PG recycling products including anhydro-muramyl tripeptide (AMP-P3) and pentapeptide (AMP-P5), along with the soluble PG precursor (UDP-M5) across different mutant backgrounds.

PG precursor levels were elevated in the Δ*dedA4* mutant (**Fig. 6A**), a phenotype which was also observed in *V. cholerae* and *S. aureus* lacking the UndP transporters Vca0040 and SAOUHSC_00846, respectively (18), suggesting DedA4 plays a role in lipid carrier recycling. Although the accumulation of PG precursors persisted in the Δ*ampD* Δ*dedA4* and Δ*ampD* Δ*dedA4* Δ*dedA5* strains, it was less pronounced than in the Δ*dedA4* single mutants (**Fig. 6A**). As expected, AMP-P3 accumulated in all Δ*ampD* derivatives, though it showed a modest decrease in the Δ*ampD* Δ*dedA4* and Δ*ampD* Δ*dedA4* Δ*dedA5* strains (**Fig. 6B**). A similar trend was observed for AMP-P5 (**Fig. 6C**). Since AMP-P5 acts an inducer and UDP-M5 as a repressor of *ampC* expression, we calculated their ratio to evaluate DedA’s role in maintaining their balance and, ultimately, its impact on beta-lactamase induction. Interestingly, the ratio of tri-and pentapeptide-AMP to UDP-MurNAc-pentapeptides dropped by 32 % in the Δ*ampD* Δ*dedA4* and 38 % in the Δ*ampD* Δ*dedA4* Δ*dedA5* strains (**Fig. 6D**) suggesting that AmpR shifted towards its repressor state. To confirm this, we measured *ampC* transcript levels using *rpsL* and *gyrB* as housekeeping controls. Expression of *ampC* was reduced 19-fold in the Δ*ampD* Δ*dedA4* double mutant, compared to the Δ*ampD* strain, which displays a strong *ampC* induction (51-fold increase relative to the wild-type) (**Fig. 7A**).

**Figure 6.**
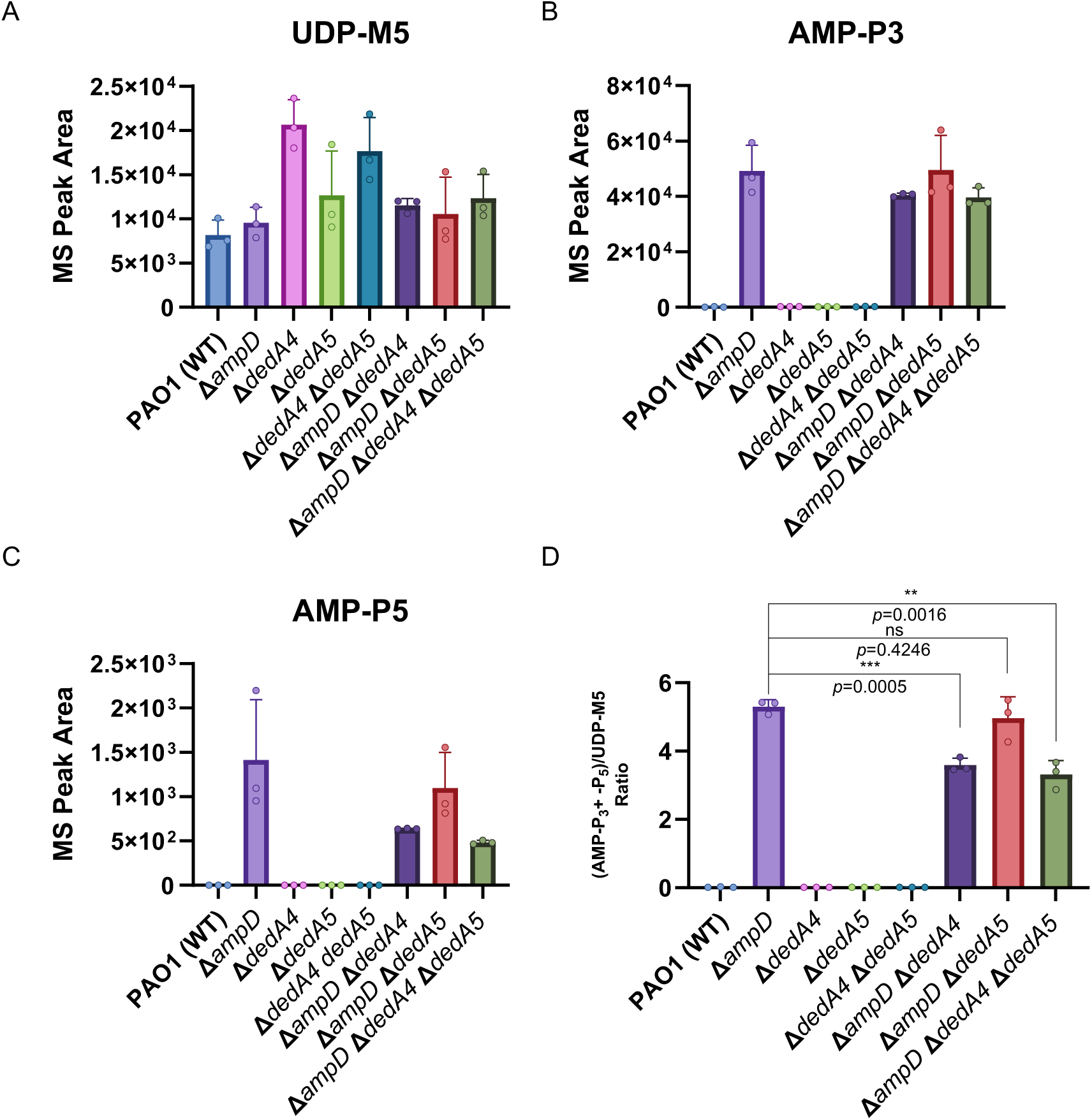
Changes in the balance of AMP-P3 + AMP-P5 / PG precursor in Δ*ampD* Δ*dedA4* confirmed by soluble muropeptide analyses. (**A-C**) Quantification of UDP-MurNAC pentapeptide (UDP-M5) (**A**), tripeptide-AMP (AMP-P3) (**B**) and pentapeptide-AMP (AMP-P5) (**C**) in strains PAO1 [WT], CF5 [Δ*ampD*], CF1842 [Δ*dedA4*], CF2034 [Δ*dedA5*], CF2037 [Δ*dedA4* Δ*dedA5*], CF1844 [Δ*ampD* Δ*dedA4*], CF2041 [Δ*ampD* Δ*dedA5*] and CF2043 [Δ*ampD* Δ*dedA4* Δ*dedA5*]. (**D**) Ratio of PG recycling products (tripeptide-AMP and pentapeptide-AMP) to PG precursors (UDP-M5) in the same strains.

**Figure 7.**
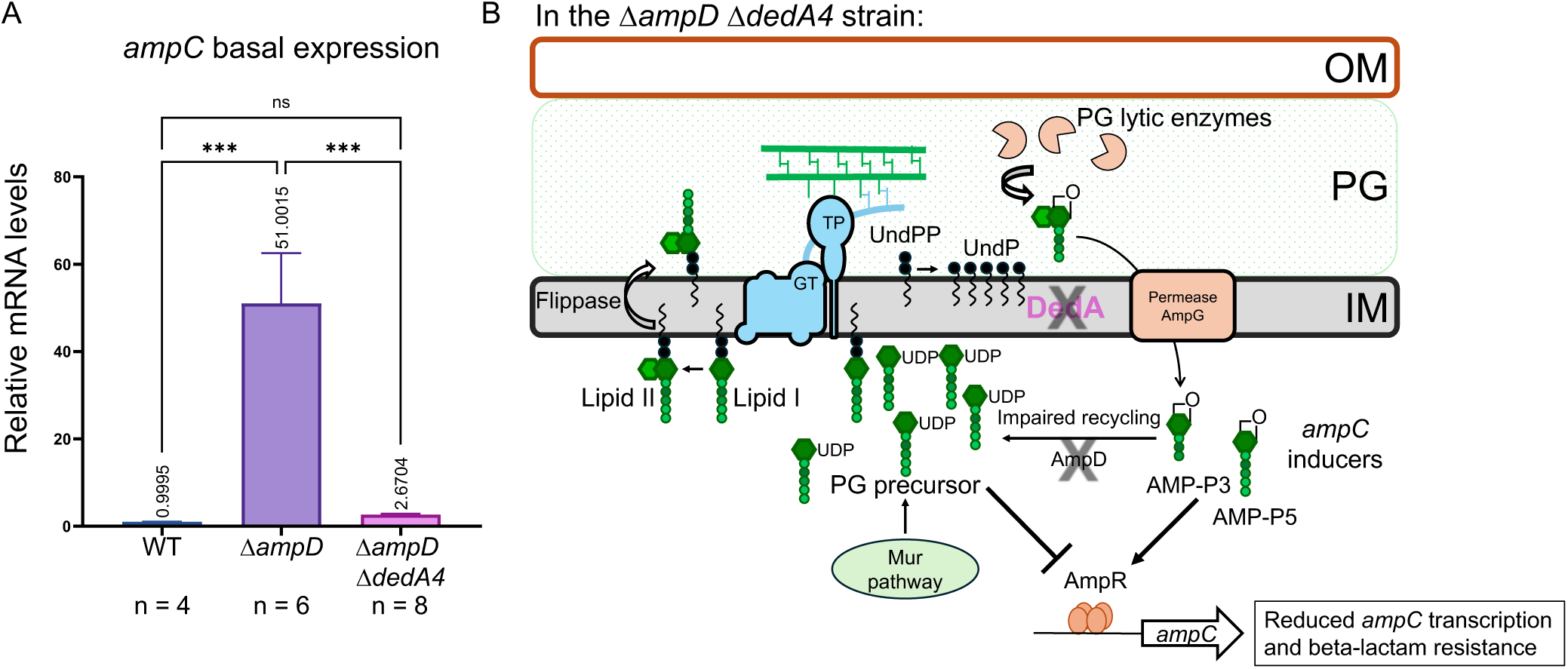
The change in AMP-P3 + AMP-P5 / UDP-M5 ratio is sufficient to switch most AmpR proteins to their repressive form. (A) Basal expression of *ampC* in PAO1 [WT], CF5 [Δ*ampD*] and CF1844 [Δ*ampD* Δ*dedA4*]. (**B**) Model illustrating the effect of *dedA4* deletion on the balance between PG recycling products and PG precursors, AmpR activity and *ampC* transcription in the Δ*ampD* background (see text for details).

Taken together, these data confirm that deletion of *dedA4* in the Δ*ampD* mutant alters the intracellular ratio of PG precursors to recycling products, shifting AmpR activity toward its repressor form. This ultimately results in lower AmpC production and restores beta-lactam susceptibility in otherwise resistant Δ*ampD* strains (**Fig. 7B**).

### Changing levels of UndP or PG precursors by expressing UppS or MurA influence the level of beta-lactam resistance

The Δ*dedA4* strain accumulates soluble PG precursors (UDP-M5) in its cytoplasm (**Fig. 6A**). We hypothesized that this accumulation may interfere with AmpR activation during cefoxitin exposure, thereby rendering the Δ*dedA4* strain more sensitive to cefoxitin than the wild-type. Wild-type *P. aeruginosa* strains are typically highly resistant to cefoxitin, as it is both a potent *ampC* inducer and an effective AmpC substrate (4). Indeed, we found that the Δ*dedA4* mutant was more susceptible to cefoxitin than both the wild-type and Δ*dedA5* strains, which display similar resistance profiles (**Fig. 8A**). We next examined whether overexpressing *uppS* (*PA3652*), which encodes a key enzyme in the *de novo* synthesis of UndP, could restore cefoxitin resistance in the Δ*dedA4* background (21). By increasing the supply of UndP lipid carriers, PG precursors can be more efficiently used for cell wall synthesis, reducing their cytoplasmic accumulation and relieving AmpR repression. As anticipated, *uppS* expression successfully restored the growth of the Δ*dedA4* strain to wild-type levels on plates containing 150 µg/ml cefoxitin (**Fig. 8B**). In contrast, increasing the intracellular concentration of UDP-M5 in a Δ*ampD* mutant, achieved by overexpressing *murA* (*PA4450*), the enzyme catalyzing the first committed step in PG precursor biosynthesis, led to heightened sensitivity to beta-lactam antibiotics (25) (**Fig. 8C**). Collectively, these findings emphasize the essential role of UndP availability in balancing PG precursors and regulating AmpC-mediated beta-lactam resistance, highlighting PG lipid carrier metabolism as a key factor in antibiotic susceptibility.

**Figure 8.**
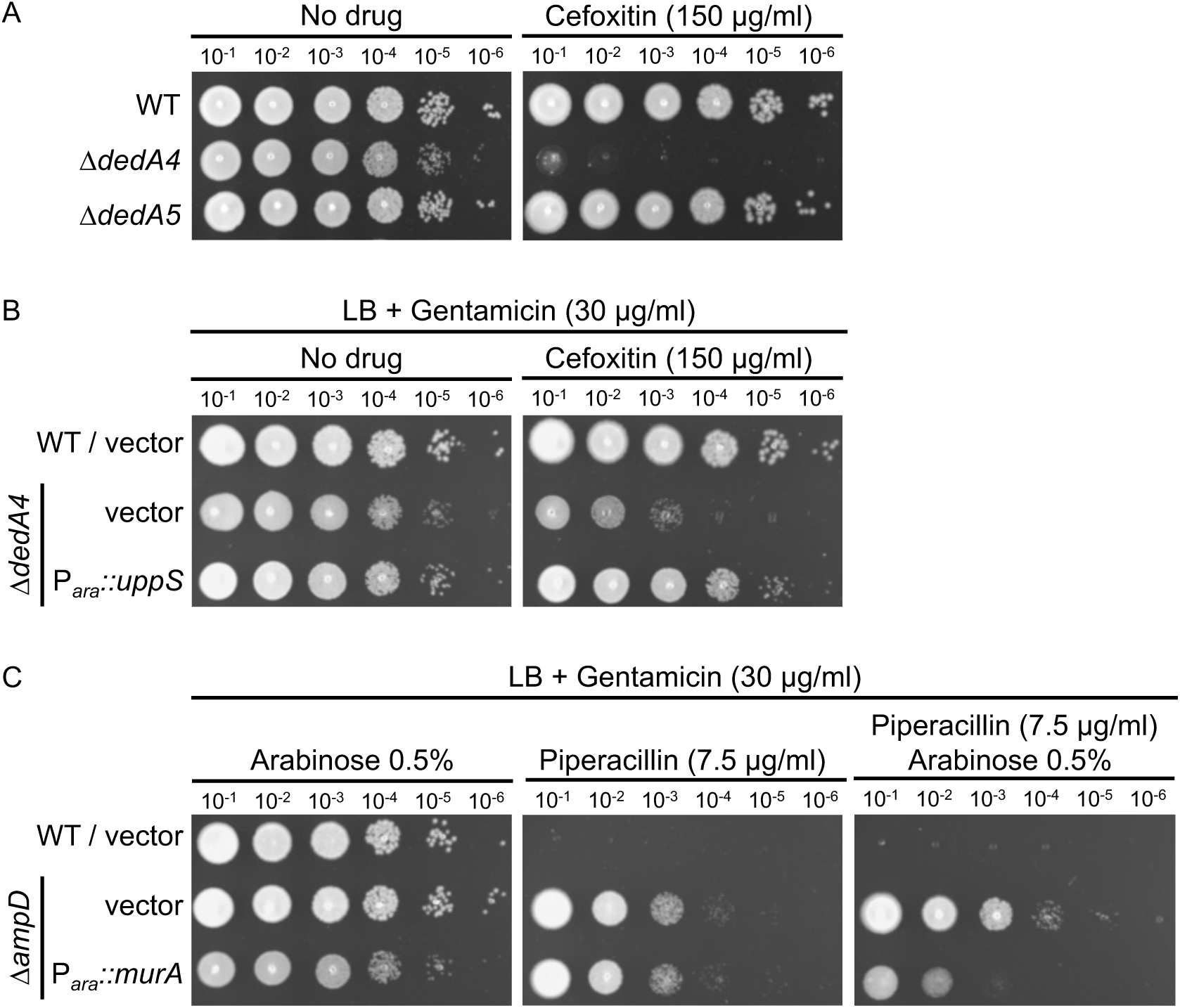
Increasing UndP or UDP-M5 levels by overproducing UppS or MurA, respectively, affects beta-lactam resistance, probably by modulating *ampC* induction. (**A**) Cultures of strain PAO1 [WT], CF1842 [Δ*dedA4*] and CF2034 [Δ*dedA5*] were serially diluted and 5 µl of each dilution was spotted onto LB agar supplemented with cefoxitin (150 µg/ml). (**B**). Cultures of strain PAO1 [WT] and CF1842 [Δ*dedA4*] with plasmids pJN105 [vector control] and pCF214 [P*_ara_*::*uppS*] were serially diluted and 5 µl of each dilution was spotted onto LB agar supplemented with gentamicin (30 µg/ml) for plasmid maintenance and cefoxitin (150 µg/ml), as indicated. (**C**) Cultures of strain PAO1 [WT] and CF5 [Δ*ampD*] with plasmids pJN105 [vector control] and pCF166 [P*_ara_*::*murA*] were serially diluted and 5 µl of each dilution was spotted onto LB agar supplemented with gentamicin (30 µg/ml) for plasmid maintenance, arabinose (0.5%) and piperacillin (7.5 µg/ml), as indicated.

## DISCUSSION

Overproduction of the beta-lactamase AmpC is a major resistance mechanism against beta-lactam antibiotics in *P. aeruginosa* and members of the Enterobacteriaceae, such as *Citrobacter freundii* and *Enterobacter cloacae* (4). In these species, loss-of-function mutations in the cytoplasmic PG recycling amidase AmpD are a common route to constitutive *ampC* expression. (26–30, 5). In such mutants, PG recycling intermediates, particularly AMP-P3, are thought to accumulate and to competitively displace the PG precursor bound to the transcriptional regulator AmpR, switching it to its activator state (12, 13).

Several strategies have been explored to circumvent AmpC-mediated beta-lactam resistance. The most established approach involves combining beta-lactams with beta-lactamase inhibitors (BLIs). While piperacillin-tazobactam was widely used against *P. aeruginosa* (31), resistance has prompted the development of newer BLI combinations, such as ceftazidime-avibactam, imipenem-relebactam, and meropenem-vaborbactam (32). These combinations improve efficacy but are still susceptible to resistance evolution (33).

Alternative strategies focus on disrupting *ampC* regulation. Inhibiting the permease AmpG prevents the uptake of muropeptides that signal AmpR activation (34, 35), while blocking the glucosaminidase NagZ interferes with the generation of strong AmpR inducers (36). Deletion of either gene or chemical inhibition (e.g., CCCP for AmpG or EtBuPUG for NagZ) reduces *ampC* expression and beta-lactam resistance in derepressed backgrounds (34–37).

Our work shows that PG precursor accumulation can be harnessed to suppress AmpC production, offering a new avenue for regulating antibiotic resistance (**Fig. 7B**). Unlike conventional strategies that inhibit inducer production, this approach indirectly drives AmpR toward its repressor state by shifting precursor dynamics, unveiling a previously overlooked mechanism with significant therapeutic promise.

To experimentally test this concept, we targeted UndP recycling to manipulate PG precursor flux. Specifically, we deleted the gene encoding *P. aeruginosa* UndP flippase, *dedA4*, in an *ampC*-derepressed Δ*ampD* background to disrupt precursor utilization and promote their cytoplasmic accumulation. This deletion prevented AmpC production, AmpC activity and beta-lactam resistance (**Fig. 2 and 3**). The UndP recycling activity of DedA4 was required to restore AmpC production in the Δ*ampD* Δ*dedA4* strain (**Fig. 4**), and heterologous complementation with the *E. coli* UndP transporters YqjA and YghB could also complement the double mutant (**Fig. 5**). We confirmed that the *dedA4* mutant had increased levels of soluble PG precursors, similar to *V. cholerae* and *S. aureus* strains deleted for DUF368 homologs (**Fig. 6A**) (18). Although PG precursor levels were not as elevated in the Δ*ampD* Δ*dedA4* strain, likely due to impaired precursor formation via the AmgK-MurU-MupP recycling pathway (38–41), the ratio of PG precursors to recycling products was significantly altered (**Fig. 6D**). This shift biased AmpR activity toward its repressor state, resulting in a 19-fold reduction in *ampC* transcript levels (**Fig. 7**). Similarly, increasing PG precursor synthesis via *murA* overexpression increases antibiotic susceptibility of the *ampD* mutant, further supporting the idea that shifting the balance towards the PG precursor interferes with AmpR activation (**Fig. 8**).

Notably, a recent study has also shown that DedA4 is required for *ampC* induction and beta-lactam resistance in a clinical bloodstream isolate of *P. aeruginosa* harboring a *dacB* mutation (PBP4 G437D) and therefore also derepressed for *ampC* (42, 43). Importantly, these results highlight that *ampC* expression depends not simply on the presence of inducers but on the ratio between PG recycling products and biosynthetic intermediates. This underscores the sensitivity of AmpR to subtle shifts in intracellular PG metabolite pools and suggests that interfering with UndP recycling is a viable strategy to manipulate *ampC* expression without targeting the enzyme or its regulators directly.

Natural products that target the UndP cycle, such as amphomycin, friulimicin B and bacitracin, bind to outward-facing UndPP or UndP, but their activity is largely limited to Gram-positive bacteria due to poor outer membrane permeability in Gram-negative species (44–46). Nonetheless, the development of smaller, membrane-permeable molecules with similar targets specificity could provide a viable strategy to reduce *ampC* expression and potentiate beta-lactam antibiotics in *P. aeruginosa* and other pathogens encoding an inducible *ampC*.

Intriguingly, other UndP-linked pathways may influence this regulatory system. For example, biosynthesis of O-antigen and capsule also depends on UndP (14). Competition between these pathways and PG synthesis could conceivably modulate precursor pools and impact AmpR signaling. Preliminary evidence from a *P. aeruginosa mucD* mutant, known to accumulate alginate (47), suggests that redirecting UndP flux toward exopolysaccharide production may also repress *ampC* induction (unpublished data). If validated, this would broaden the metabolic processes implicated in beta-lactam resistance and support the idea that PG precursor homeostasis is a central regulatory hub. We aim to explore this in future work.

In summary, our findings reveal that AmpC expression is controlled by a finely tuned balance between PG precursors and recycling products. Disruption of UndP recycling shifts this balance toward precursor accumulation, thereby repressing AmpR and *ampC* even in genetically derepressed backgrounds. This work identifies a novel strategy to suppress beta-lactam resistance and underscores the importance of lipid carrier dynamics in coordinating antibiotic response pathways. Notably, similar links between membrane homeostasis and resistance have been observed in *B. subtilis* and *Burkholderia thailandensis*, where mutations in *dedA* homologs modulate susceptibility to duramycin and colistin, respectively (24, 48). Targeting UndP recycling offers a promising new direction for adjunctive therapies that could resensitize AmpC-producing bacteria to existing beta-lactam antibiotics.

## METHODS AND MATERIALS

### Media, bacterial strains, and plasmids

*P. aeruginosa* PAO1 cells were grown in LB (1% tryptone, 0.5% yeast extract, 1% NaCl). As indicated, the medium was supplemented with 1 mM IPTG (isopropyl beta-D-1-thiogalactopyranoside) or 0.5% arabinose. For plasmid maintenance, gentamicin (Gm) was used at a concentration of 30 μg/mL. Unless otherwise indicated, antipseudomonal antibiotics for viability/sensitivity assays were used at 3, 5, 7.5 or 10 μg/mL (piperacillin; Pip or ceftazidime; Caz), as indicated. All *P. aeruginosa* strains used in the reported experiments are derivatives of PAO1. *E. coli* cells were grown in LB. For plasmid maintenance or selection in *E. coli*, antibiotic concentration used was 15 μg/mL (gentamicin; Gm). The bacterial strains and plasmids used in this study are listed in **Table S1, S2 and S3**. Detailed descriptions of the strain and plasmid construction procedures can be found in the Supplementary Material.

### *P. aeruginosa* viability assay

For viability assays, overnight cell cultures were normalized to an OD_600_ of 0.1 and subjected to serial 10-fold dilutions with LB. Five microliters of each dilution was then spotted onto the indicated agar and plates were incubated at 30°C for 24 hours prior to imaging.

### *P. aeruginosa* electroporation

*P. aeruginosa* strains were made competent using previously described methods (49). Briefly, 4 mL of overnight cultures grown at 37°C were centrifuged and washed twice with 1 mL 300 mM sucrose. Cell pellets were resuspended in 500 µL of 300 mM sucrose and 100 µL were used for electroporation. 1µL of replicative plasmid was used for the electroporation (Gene Pulser, Bio-Rad), using the following settings: 25 mF, 200 O, 2.5 kV. LB medium (1mL) was added and the cells incubated with shaking (200 rpm) for 1 h at 37 °C. Cells were then plated on the appropriate selective medium.

### AmpC beta-lactamase activity assay

AmpC activity was assessed using nitrocefin hydrolysis. Overnight bacterial cultures were subcultured 1:20 in 3 mL LB and grown for 4 hr at 30°C and 200 rpm. Following incubation, 1 mL of culture was pelleted at 5’000 x g for 5 minutes, washed once with 1 mL of 50 mM sodium phosphate buffer (pH 7.0) and resuspended in 2 mL of the same cold buffer. The samples were frozen on dry ice for 15 minutes and thawed on ice. Samples were placed on ice and lysed at 4°C by sonication with a microprobe (Q800R2, QSonica, Newtown, Connecticut, USA). Sonicated samples were centrifuged at 12,000 x g for 10 minutes at 4°C and supernatants were collected. The protein concentration was determined using a Bradford assay (50) with bovine serum albumin (BSA) as the standard (G-Biosciences, Geno technology inc., Saint-Louis, Missouri, USA). Nitrocefin hydrolysis assays were performed in 96-well plates. Each reaction had a final volume of 250 μl of 50 mM sodium phosphate buffer (pH 7.0) containing 10 μg of protein and 20 μg of nitrocefin (Thermo Fischer Scientific™ Oxoid, Waltham, Massachusetts, USA). Nitrocefin hydrolysis was monitored by measuring the absorbance at 340 and 486 nm every 2 minutes for 120 minutes at 30°C.

### Antibiotic sensitivity assays

Antibiotic sensitivity assays were performed using broth microdilutions. Overnight cell cultures were normalized to OD_600_ of 0.0005 in LB and the indicated concentrations of piperacillin, ceftazidime, A22, tobramycin or gentamicin, and grown for 24 hours at 30°C prior to taking optical density readings (Biotek Epoch 2, Agilent, Santa Clara, California, USA). Broth microdilution MIC assays were performed three or four times independently, each with three technical replicates.

### Intracellular Soluble muropeptide analysis

To determine the presence and levels of intracellular soluble muropeptides, bacteria were grown until late exponential phase (roughly OD_600_ 0.7) in LB media before being cooled on ice for 10 min and normalized to the same OD_600_. Cells were then harvested by centrifugation at 10,000 × *g* for 10 min. The supernatant was discarded, and the cell pellet was washed three times in ice-cold 0.9% NaCl, resuspended in 0.9% NaCl so that the cells are 20 times concentrated and boiled for 10 min before centrifugation at maximum speed in a benchtop centrifuge for 10 min to remove the proteins and insoluble fraction. The supernatant was used for further analysis by LC-MS.

### Peptidoglycan isolation

Cells from 0.2 L cultures of overnight stationary phase were pelleted at 5,250 x g and resuspended in 5 ml of PBS, added to an equal volume of 10% SDS in a boiling water bath and vigorously stirred for 3 h, then stirred overnight at room temperature. The insoluble fraction (peptidoglycan) was pelleted at 400,000 x g, 15 min, 30 °C (TLA-100.3 rotor; OptimaTM Max ultracentrifuge, Beckman) and resuspended in Milli-Q water. This step was repeated 4-5 times until the SDS was washed out. Next, peptidoglycan was treated with proteinase K (40 μg/ml in 100 323 mM TrisHCl pH 8.0), 30 min at 37 °C and then boiled in 1% SDS for 2 h to stop the reaction. After SDS was removed as described previously, peptidoglycan samples were resuspended in 200 μL of 50 mM sodium phosphate buffer pH 4.9 and digested overnight with 30 μg/ml muramidase (from *Streptomyces albus*) at 37 °C. Muramidase digestion was stopped by heat-inactivation (boiling during 5 min). Coagulated protein was removed by centrifugation (20,000 x g, 15 min). The supernatants (soluble muropeptides) were subjected to sample reduction. First, pH was adjusted to 8.5-9 by addition of borate buffer 0.5 M pH 9 and then muramic acid residues were reduced by sodium borohydride treatment (NaBH_4_ 10 mg/ml final concentration) during 30 min at room temperature. Finally, pH was adjusted to 2.0-4.0 with orthophosphoric acid 25% prior to analysis by LC.

### LC-MS analysis

Chromatographic analyses of muropeptides were performed by Ultra Performance Liquid Chromatography (UPLC) on an UPLC system (Waters) equipped with a trapping cartridge precolumn (SecurityGuard ULTRA Cartridge UHPLC C18 2.1 mm, Phenomenex) and an analytical column BEH C18 column (130 Å, 1.7 μm, 2.1 mm, Waters) maintained at 45 °C. Muropeptides were detected by measuring the absorbance at 204 nm using an ACQUITY UPLC UV−visible Detector. Muropeptides were separated using a linear gradient from buffer A (Water + 0.1 % (v/v) formic acid) to buffer B (Acetonitrile 100 % (v/v) + 0.1 % (v/v) formic acid) over 15 min with a flowrate of 0.25 ml/min. The QTOF instrument was operated in positive ion mode, with data collection performed in untargeted MS^e^ mode. The parameters were set as follows: capillary voltage 3.0 kV, source temperature 120 °C, desolvation temperature 350 °C, sample cone voltage 40 V, cone gas flow 100 L h^−1^ and desolvation gas flow 500 L h^−1^. Mass spectra were acquired at a speed of 0.25 s/scan. The scan was in a range of 100–2000 m/z. Data acquisition and processing was performed using MassLynx or UNIFI software package (Waters Corp.). The quantification of muropeptides was based on their relative abundances (relative area of the corresponding peak) and relative molar abundances. A table of all the identified muropeptides and the observed ions is provided (**Table S4**).

### Statistical Analysis and Reproducibility

Statistical evaluations were conducted using Prism 8.0 (GraphPad Software, USA). For comparisons, two-tailed unpaired t-tests were applied. A p-value threshold of less than 0.05 was considered statistically significant, with significance levels denoted as follows: *p* < 0.05, *p* < 0.01, \**p* < 0.001, and \*\**p* < 0.0001. All assays incorporated appropriate control groups, and both experimental and control conditions were carried out using isogenic strains. Experiments included a minimum of three independent biological replicates.

### Immunoblotting

Overnight bacterial cultures were subcultured 1:100 in 3 mL of appropriate medium and grown for 4 hours. Bacteria were normalized to an OD_600nm_ of 0.5 in 1 ml, pelleted and resuspended in a final volume of 100 µl. 20 µl of Laemmli buffer were added and the samples were then boiled for 10 minutes at 95°C. Immunoblotting was performed by first separating 10 µL of each sample on 12% SDS-PAGE (<u>p</u>oly<u>a</u>crylamide <u>g</u>el <u>e</u>lectrophoresis) gels at 70V for 30 minutes and 120V for an hour. Proteins were transferred at 90V for an hour to a 0.2 μm PolyVinylidene DiFluoride (PVDF) membranes (Immobilon-FL PVDF membrane, IPFL00010, Merck) previously soaked in methanol and rinsed with transfer buffer. Membranes were stained with Ponceau S solution to ensure equal loading in each lane and transfer. Membranes were blocked using 5% (w/v) skim milk in Tris-Buffered Saline (10 mM Tris-HCl pH 7.5, 150 mM NaCl) supplemented with 0.1% (v/v) Tween-20 (TBS-T) for 1 hour. Membranes were incubated for 1 hour with α-AmpC primary antibody (1:1000 dilution in 5% skim milk in TBS-T, MyBioSource, MBS1493275, San Diego, USA) or α-FLAG primary antibody (1:1000 dilution in 5% skim milk in TBS-T, F7425, Merck). The membranes were washed four times in TBS-T for 5 min each before incubation for 1 h with secondary antibody (anti-rabbit IgG HRP, 1: 10000 dilution, A0545, Merck) in TBS-T with 5% (w/v) skim milk powder. The membranes were then washed four times with TBS-T for 5 min each before developing using Immobilon Western Chemiluminescent HRP Substrate (WBKLS0100, Merck) and imaged using the FUSION FX Spectra imaging platform (VILBER). Images were acquired with the Evolution Cap Edge software.

### RT-qPCR

The relative expression level of *ampC* was determined by quantitative reverse transcription PCR (RT-qPCR). Total RNA was isolated from 1 ml of bacterial cultures in the logarithmic growth phase using the RNeasy 96 QIAcube HT Kit (Qiagen) according to the manufacturer’s instructions. One microgram of total RNA from each sample was reverse transcribed using the QuantiTect® Reverse Transcription Kit (Qiagen) with random hexamers. The resulting cDNA was diluted 1:5 and analyzed in duplicate for quantitative PCR with SYBR® Green master mix (BIO-RAD) using the QuantStudio™ 3 Real-Time PCR System (Applied Biosystems).

The threshold cycle (Ct) for each sample, corresponding to the PCR cycle at which fluorescence exceeds a predefined threshold, was determined using the QuantStudio™ software. The *rpsL* and *gyrB* housekeeping genes were used as internal controls. The primers used are listed in **Table S5**. The mean values of relative mRNA expression (calculated as 2^(−ΔΔC_T_ )^(51)) from two independent biological replicates and two technical replicates were considered for analysis. The wild-type strain was used as the control.

Primers efficiencies were assessed by generating a standard curve with serial dilutions of a cDNA template. Primers efficiencies were determined to be ≥ 90%, with no more than a 5% variation between each pair of primers. A final melting curve step was systematically included to verify the specificity of the amplification.

## ACKNOWLEDGEMENTS

The authors would like to thank all members of Cava and Fumeaux labs for advice and helpful discussions. Research in the Fumeaux lab is supported by a PRIMA grant from the Swiss National Science Foundation (#PR00P3_185713), a grant from the Swiss Cystic Fibrosis Foundation (CFCH), the Pierre Mercier Foundation for Science and the Jubiläumsstiftung von Swiss Life. Research in the Cava lab is supported by The Swedish Research Council (VR), The Knut and Alice Wallenberg Foundation (KAW), The Laboratory of Molecular Infection Medicine Sweden (MIMS), Cancerfonden and The Kempe Foundation.

